# ASTA-P: a pipeline for the detection, quantification and statistical analysis of complex alternative splicing events

**DOI:** 10.1101/2023.08.28.555224

**Authors:** Kanupriya Tiwari, Lars Keld Nielsen

## Abstract

Alternative splicing dramatically increases the repertoire of the human transcriptome and plays a critical role in cellular differentiation. Long read sequencing has dramatically improved our ability to explore isoform diversity directly. However, short read sequencing still provides advantages in terms of sequencing depth at low cost, which is important in comparative quantitative studies. Here, we present a pipeline called ASTA-P for profiling, quantification, and differential splicing analysis of tissue-specific, arbitrarily complex alternative splicing patterns. We discover novel events by supplementing existing annotation with reconstructed transcripts and use spliced RNA-seq reads to quantify splicing changes accurately based on their unique assignments. We used simulated RNA-seq data to demonstrate that ASTA-P provides a good trade-off between discovery and accuracy compared with several popular methods. Further, we applied ASTA-P to analyse AS patterns in real data from hiPSC derived cranial neural crest cells capturing the transition from primary neural cells into migratory cranial neural crest cells, differentiated by their expression of the transcription factor, SOX10. Our analysis revealed a significant splicing complexity, i.e., numerous AS events that cannot be described using the conventionally analysed 2D splicing event patterns. Such events are misclassified when analysed using current differential splicing analysis methods. Thus, ASTA-P provides a new approach for studying both conventional and complex splicing across different cellular conditions and the dynamic regulation of AS. The pipeline is available at https://github.com/uqktiwar/ASTAP/tree/main

## 1. Introduction

Alternative splicing (AS) is an important regulatory mechanism for modulating gene expression or protein function across different biological conditions. Vast amounts of RNA-seq data surveying different physiological and disease conditions are continuously being generated and analysed to advance our knowledge of AS mechanisms, and it is now estimated that more than 95% of human genes express alternative isoforms [1]. However, due to the lack of supporting proteomic evidence, exactly how many of these novel isoforms are functional is a matter of ongoing debate [2]. Transcript sequence variations due to alternative splicing have been analysed using a variety of data sources including cDNA, ESTs, and protein domain sequences mapped to the genome, as well as, by aggregating expression information from tiling and exon-exon junction microarrays [3]. These analyses involved pairwise comparisons of say, an EST sequence to either a single reference transcript or by conducting exhaustive pairwise comparisons against all known isoforms of a gene [3], [4]. Such pairwise analyses uncovered the set of commonly analysed “classical” binary splicing patterns, namely: cassette exon skipping, alternative 5’ or 3’ splice site (SS) usage, mutually exclusive exons, and intron retention. However, overlapping instances of these patterns were often observed, indicating that a more complex cumulative AS structure would be a better descriptor of the splicing variation observed for some genes [3]. Consequently, a few methods conducting a more generalized search of AS patterns emerged. For instance, Nagasaki et al. (2005) and Sammeth et al. (2007) respectively developed the ASTI [5] and ASTALAVISTA [4], [6] algorithms to survey the different array of splicing patterns observed in the human transcriptome using mRNA and EST evidence. Both studies detected a greater variety of patterns beyond the classical patterns. Nagasaki et al. used 2D bit array representations of 65,041 Unigene mRNAs mapped to the human genome to identify 124 distinct types (11,489 total) of “AS units”, defined as combinations of variably expressed genomic regions flanked by common exonic or intronic regions [5]. On the other hand, Sammeth et al., clustered GTF formatted mRNA and EST sequences to construct alternative splicing graphs (ASG). The ASGs were then parsed to mine arbitrarily complex AS events as “bubbles” in this ASG. Due to their use of ESTs to supplement mRNA and annotated RefSeq evidence, they uncovered more pervasive AS, retrieving >90,000 AS events, more than a quarter of which (27%) were complex events producing more than two variants [4].

Advances in RNA-sequencing technologies has dramatically expanded our ability to survey transcriptomes, and dozens of methods offering not just AS detection but also quantification and differential analysis capabilities have been developed. Among these, popular event-based tools, such as rMATS [7], have been used to make valuable contributions to the field through the detection of key differential splicing events between different conditions, as well as, understanding sequence-based regulation of different AS types. However, their analysis is limited to the detection of classical binary patterns and they often predict a significant fraction of overlapping events. Only very recently, methods that survey splicing patterns from RNA-seq data in a generalised manner have emerged. For instance, Vaquero-Garcia et al. developed the tool, MAJIQ, which detects “local splicing variations” (LSVs) involving two or more splice junctions that share a common source (5’) or sink (3’) splice site. They uncovered that complex splicing variations (>= 3 junctions with a shared site) occur at a greater frequency than previously appreciated, accounting for 30% of all LSVs detected in human genes [8]. Another recently published tool, JUM, defines AS structures as clusters of splice junctions akin to MAJIQ’s LSVs and is similarly able to detect complex combinations of the binary patterns [9]. Blencowe et al. published Whippet, which combines annotation information and RNA-seq data to construct denovo contiguous splice graphs (CSGs). Within these graphs, non-overlapping exonic segments represent nodes and, directed edges connect the nodes in each unique path through the CSG. AS events are defined on a per-node basis as the set of all paths in the CSG skipping or including a node and the complexity of an event is measured as K(n) = log_2_(n = number of paths defining the event) [10].

In this paper, we present a pipeline, ASTA-P (Figure 1), for the analysis of arbitrarily complex splice patterns, using ASTALAVISTA [6] to mine complete splicing events of different dimensions, followed by quantification with a custom script, and modelling the event counts using the Dirichlet-multinomial regression framework implemented in the R-package, DRIMseq [11]. We combine full-length transcript reconstruction for enriching the existing annotation model before assembling the splicing graph for each gene. We compare our pipeline against four other event-based methods - two commonly used classical event analysis methods (rMATS [7] and SUPPA [12]) and two of the recently published complex-splicing analysis tools (Whippet [10] and JUM [9]) - w.r.t to their ability to detect differential splicing. In addition to these tools, we also included a Salmon + DRIMseq [11], [13] pipeline in the comparison, since DRIMseq was originally developed for DS analysis by modelling full-length transcript counts (or exon-level counts akin to DEXseq [14]). We used simulated data to examine the effects of varying the degree of differential splicing, sequencing depth, and incomplete annotation on the detection of differentially spliced genes by these tools. Further, our analysis of real RNA-seq data from cranial neural crest cells (CNCCs) revealed a significant occurrence of complex splicing patterns. We also evaluated some of the features associated with these events as compared to the classical 2D patterns.

**Figure 1.**
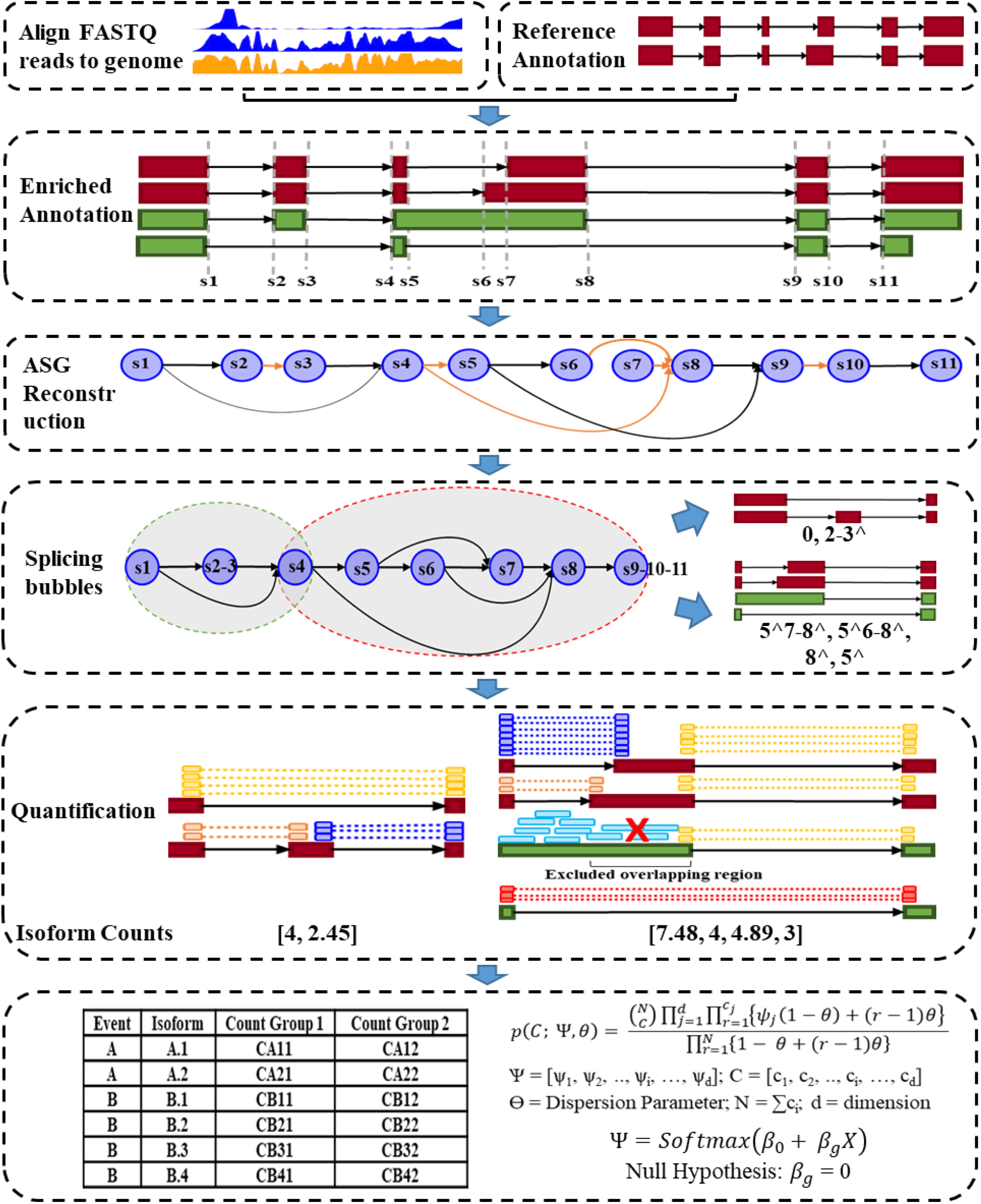
The ASTA-P pipeline for detection, quantification and statistical analysis of complex AS events by combining published tools with a custom quantification method. We employ a hybrid approach combining full-length transcript reconstruction for enriching the existing annotation model and assembling a splice graph for each gene. This is followed by mining and quantification of local splice events including binary as well as high dimensional patterns from the splice graph. Once we have obtained the splice events and RNA-seq read counts corresponding to different isoforms generated from each event, we model the isoform proportions using Dirichlet-multinomial regression to assign proportions and identify differentially spliced events.

## 2. Results

### 2.1 The ASTA-P pipeline

#### 2.1.1 Splicing Event Detection as Bubbles in the Alternative Splicing Graph

We use alternative splicing graphs (ASGs) as the basic units to identify splicing events. The alternative splicing graph (ASG = (V, E)) of a gene (G) is a directed acyclic graph with nodes corresponding to transcript starts, ends and splice sites, and edges corresponding to exonic regions and splice junctions, directed from the 5’ to the 3’ end [6], [15]. The set of transcripts detected in an RNA sample is provided to ASTALAVISTA for splicing graph construction and mining arbitrarily complex splicing events. In order to identify novel splicing events, RNA-seq data are aligned to a reference genome and the alignments used for transcript reconstruction to supplement the existing annotation. Subsequently, all transcripts (T) of a gene described in the gene transfer format (GTF) file are combined to construct the ASG.

ASTALAVISTA detects splicing events as “bubbles” in the ASG [4]. Each bubble is an induced subgraph of the ASG, specified by a set of flanking constant nodes (splice sites), and a sequence of internal (variable) nodes or splice chains describing each isoform generated from the event. Briefly, each node (s_i_ ϵ V) in the ASG represents a genomic site characterised by a tuple of attributes: {pos(s_i_), type(s_i_), support(s_i_)}. For any site (s_i_), pos(s_i_) gives its absolute genomic position, type(s_i_) indicates whether the site is an internal donor/acceptor splice site or a transcript start/end site. The transcripts containing s_i_ as a sequence boundary form the support(s_i_) set for the site. Directed edges (e_k_: s_i_ -> s_j_ ϵ E) between pairs of nodes ((s_i_, s_j_): pos(s_i_) < pos(s_j_)) represent exonic or intronic sequences characterised by the tuple – {type(e_k_), support(e_k_)}. For any edge (e_k_: s_i_ -> s_j_ ϵ E), type(e_k_) indicates whether it is an intronic or exonic sequence supported by the set of transcripts - support(e_k_).

Only edges with non-empty support are included in the ASG. A variant (p), highlighted in **Figure 2**, represents a path between any pair of nodes (s_m_, s_n_: pos(s_m_) < pos(s_n_)) in the ASG supported by a non-empty set of transcripts Xp (≠ F). A **complete AS event** is defined as the maximal set of overlapping variants (P) that share common end-points (s_m_, s_n_) and at least one intervening site (s_i_: pos(s_m_) < pos(s_i_) < pos(s_n_)) in the variant subgraph is of type(s_i_) ϵ {Donor, Acceptor}. Variant sets that are not delimited by common internal sites but by transcript start or end sites describe alternate transcription start/end events, which we are not analysing in this work.

**Figure 2.**
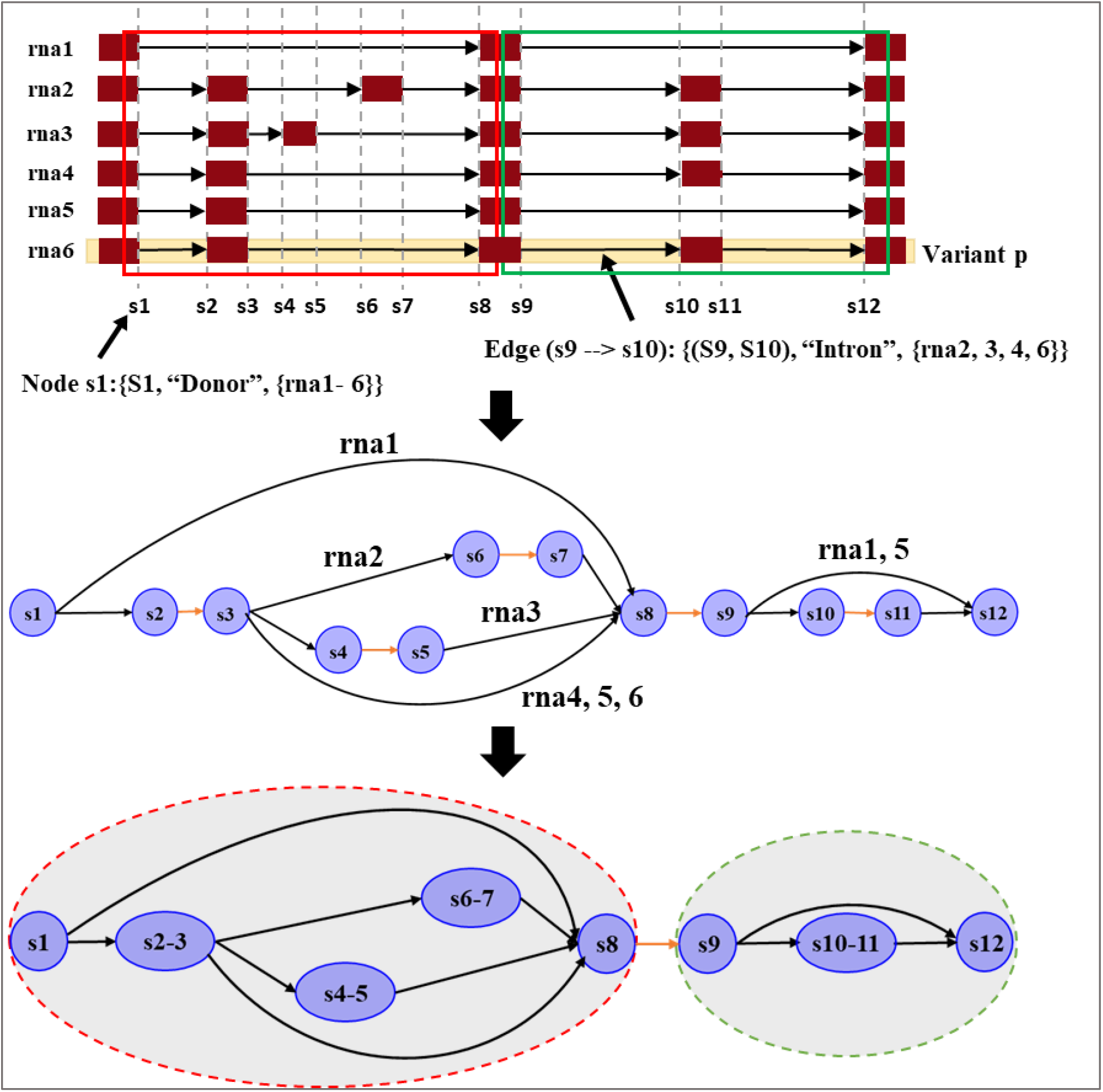
ASTALAVISTA mines complete AS events as bubbles in a splicing graph.

In order to mine complete AS events, ASTALAVISTA appends a root node (R: {-∞, root, T}) and a leaf node (L: {+∞, leaf, T}) at the ends of the ASG by extending edges from all transcript start/end sites to the root/ leaf respectively. They collapse all non-informative vertices (u) with in-degree(u) = out-degree(u) = 1. A “**bubble**” (s, t, χ_st_) is then defined as the subgraph of the ASG delimited by a pair of vertices s and t (pos(s) < pos(t)) that contains at least two variants. Vertex s is the ‘‘source’’ and t is the ‘‘sink’’ of the bubble. In order to find bubbles containing alternative splicing events of any dimension d, the algorithm iterates over all nodes with in-degree >= d in genomic order as potential sinks (t ϵ V). For each sink (t), all preceding nodes (s ϵ V: pos(s) < pos(t) & out-degree(s) >= 2), in reverse genomic order, are examined until the first node with support(s) ⊇ support(t) is found. Once the ends (s, t) of a bubble have been identified in this way, the set of all transcripts supporting t is iteratively partitioned by intersecting it with the transcript support of all edges between s and t one by one resulting in the ‘‘partition set’’ χ_st_. Once the size of the partition set |χ_st_| >= d, all d-tuples in χ_st_ represent d-dimensional AS events. The reader is referred to [4] for a detailed description of the search algorithm. The complete AS events mined as such comprise the classically analysed AS patterns (alternative 5′ splice site, A5SS; alternative 3′ splice site, A3SS; skipped cassette exon, SE; mutually exclusive exon, MXE; intron retention, IR) as well as non-classical patterns, such as multi-exon-skipping events and high dimensional events generating > 2 isoforms. For example, **Figure 8C** depicts an example of a 5D AS event in the transcripts from the *PRSS56* gene detected in the cranial neural crest cells delamination dataset.

#### 2.1.2 Splicing Event Quantification

Before quantification, we classify the events into those generating at least one intron retaining (IR) isoform (IR events) or no IR isoform (Not-IR events). Individual isoforms from these events can be defined by a unique combination of quantifiable “features”. These features comprise only splice junctions for Not-IR events and a combination of junctions and retained intronic regions for the IR events. For Not-IR events, we count splice junction spanning or “split” read alignments to quantify the underlying isoforms. For multi-mapping reads, the contribution towards any feature count is adjusted by the number of alignments. In order to normalize the count along the length of an isoform, the final count is determined by taking the geometric mean of the counts for all features. For isoforms containing features with zero counts, a pseudo count of 1 is added to that feature before taking the geometric mean. For IR events, in addition to the reads spanning any splice junctions, we also consider un-spliced reads that align within the retained intron. Often retained introns overlap exonic regions in other isoforms, which results in intermittent regions of high counts within or at the edges of the intron simply by way of such overlap. In order to avoid counting such erroneously amplified IR regions, we first extract unique regions for each retained intron. Then, the contribution of a read towards the final count is only considered if it either aligns completely within or extends into these unique regions. Counts for the different features are combined as for Not-IR events using the geometric mean to get the final isoform-level counts. Our counting method relies on junction counts (and intronic region counts for IR events) and with increasing dimensionality, the fractions of junctions shared between different isoforms of an event increases. As a result, lowly expressed isoforms may be assigned reads due to the presence of shared junctions with higher expression/dominant isoforms. Therefore, we enforced feature level count thresholds to filter out the lowly expressed isoforms. For an isoform with unique features, we require at-least one unique feature to be expressed with a threshold number (N_th_) of uniquely mapped reads across a certain number of samples (N_s_). Isoforms without any unique features are retained if all their junctions are expressed in accordance with the (N_th_, Ns) parameters. The value of N_th_ and N_s_ are set based on the number of replicates and depths of the RNA-seq samples being analysed. In some cases, once the lowly expressed isoforms are removed, due to a reduction in variability of the internal structure, the isoforms are no longer delimited by the original set of flanking sites, and we re-estimate the common flanks and splice chains using the remaining isoforms. After this, we further filter the events to remove low dimensional events that are fully contained within any predicted HD events to come up with a final set of events for the next step – i.e. modelling isoform proportions.

#### 2.1.3 Statistical Modelling of Splicing Dynamics

Currently, the most commonly used distribution for modelling RNA seq counts is the Negative Binomial distribution (NB)[16]–[19]. It is a natural extension of the single parameter Poisson distribution and accounts for additional experimental and measurement variation through a dispersion parameter. NB based generalized linear model (GLM) regression is almost ubiquitously applied for conducting gene level differential analyses. However, at the transcript-level, treating each transcript’s counts as univariate Poisson/NB distributed with independent mean parameters ignores the correlated structure of this data and only reveals differential transcript expression between conditions rather than differential transcript usage. A categorical (binomial or multinomial) distribution is more suitable for modelling the proportion of total reads generated by each transcript of a gene and a significant change in these proportions for the transcripts reveals differential splicing between the conditions tested. Additionally, ordinary categorical models can be extended to account for over-dispersion using different approaches including modifying the variance specification (quasi-binomial models) or by employing a hierarchical modelling approach as in rMATS[7]. Within ASTA-P, we employ a hierarchical modelling approach. For each event, the final proportions for the isoforms are modelled using the Dirichlet-multinomial distribution[20], [21]. According to the DM framework, the read counts (C = [c1, c2, .., ci, …, cd]) assigned to the isoforms generated through a d-dimensional AS event vary according to a multinomial distribution with parameters (N, Ψ).

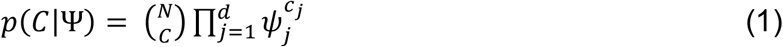

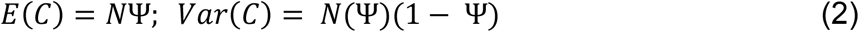

In addition, the isoform proportions (Ψ) themselves vary according to a Dirichlet distribution.

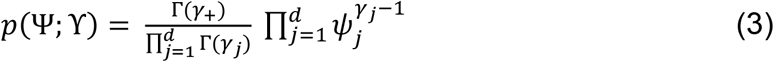

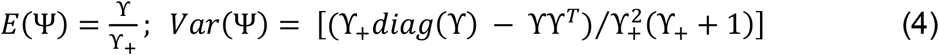

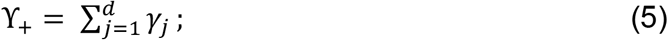

The final form of the DM distribution is achieved by integrating the product of the multinomial and Dirichlet distribution over a d-dimensional simplex and transforming the parameters as shown below.

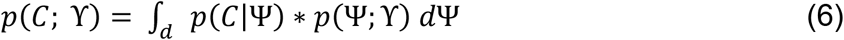

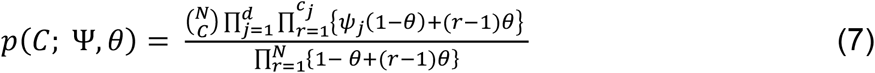

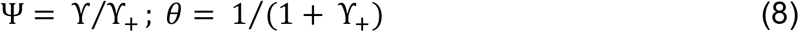

Here, θ represents the dispersion parameter used to model the over-dispersion in the isoform counts. We use the Dirichlet Multinomial (DM) regression implemented in the DRIMSeq R-package [20] to model the event counts as a function of the experimental covariates. For single condition experiments, we obtain the final proportions by specifying an intercept only model. For multi-condition experiments, we regress the mean parameters against a non-linear function of the experimental design covariates (X).

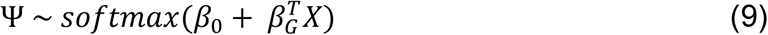

The likelihood ratio (LRT) of the full model vs a null model varies asymptotically as a χ^2^ distribution with dof = (d-1) x (t-1) and is used to test the null hypothesis that isoform proportions do not vary significantly between experimental groups (β_G_ = 0).

### 2.2 RNA-seq simulations and Performance Metrics

We evaluated six differential splicing (DS) detection methods using simulated data generated according to the workflow described in [22]. The DS analysis was limited to a control versus treatment comparison with gene-level counts for each condition sampled from NB distributions generated using a real SOX10^−^ (control) and SOX10^+^ (treatment) neural crest datasets (see Materials and Methods). Of all genes identified as multi-transcripts genes in the original dataset, 2000 were randomly selected as “true DS genes” for simulating differential splicing (**Table S1**). Of these, 849 were the result of internal AS events, whereas the remainder represent alternative transcription start or termination events. In order to simulate different degrees of DS, the PALT parameter was set to 0.2 for the control group and 0.4 or 0.6 for the treatment groups, corresponding to moderate and high degree of DS, respectively. Using FluxSimulator [23], we conducted 9 simulations to produce stranded, 100bp paired-end, triplicate RNA-seq samples for each group (control+PALT0.2, treat+PALT0.4, treat+PALT0.6) at three depths (15M, 30M, and 60M), resulting in a total of 27 fastq files. Even for paired-end simulations FluxSimulator produced single fastq files containing both reads, which we further parsed to generate separate files containing the forward and reverse reads. These reads were then mapped onto the hg19 human genome build using STARv2.6 [24] to generate BAM files required for running ASTA-P, rMATs, and JUM. Since rMATS and ASTA-P only survey internal splicing events, we evaluated the performance of all tools using only the subset of 849 genes with AS.

The tools were evaluated by visualizing their receiver operating characteristic (ROC) curves and computing the area under the curves (AUCs). For a binary classification task, such as this one, the ROC curve depicts the relationship between the true positive rate (TPR) and false positive rate (FPR) at different score thresholds used for classifying cases into one category vs another. The AUC score quantifies the probability that the classifier will assign a higher score to a positive case as compared to a negative case. Further, we also evaluated the actual FDR (1 - precision) and recall for each tool at three FDR thresholds (0.01, 0.05 and 0.1) that are commonly used to select significantly differentially spliced genes. While the ROCs illustrate the ability of a method to classify both positive and negative classes correctly, the FDR-Recall curves allow us to focus in on the performance for the positive (DS genes) class, which is especially important for imbalanced classification problems such as this one with an abundance of negative instances (10,941) as compared to positive instances (849). We parsed the output of the evaluated tools, to compile a list of genes, their associated probabilities of DS, and the maximum absolute change in proportion (ΔPSI) between the two conditions. The probability of DS was computed as 1-Padj for ASTA-P, rMATS, and JUM. For Whippet, the DS posterior probability that it computes for each event was used *as is*.

At any probability cut-off, the TPR or recall denotes the percentage of correctly called DS genes among all tested true DS genes, and the FPR denotes the percentage of incorrectly called DS genes among all tested true non-DS genes. The FDR denotes the fraction of correctly called true DS genes among all predicted DS genes by each tool. Furthermore, in addition to statistical testing, a commonly followed practice in the differential splicing field is to apply an additional |ΔPSI| cut-off for deciding the final set of differentially spliced events. This practice improves the accuracy of detection by filtering out statistically significant but low (< 10%), likely false positive changes in splicing. We incorporated this into our evaluation and computed two sets of evaluation metrics – “raw metrics” wherein genes were classified solely based on the probability cut-off and “delta metrics” wherein a gene was classified as DS only if the associated |ΔPSI| >= 10%, in addition to the probability cut-off. The ΔPSI threshold for DS genes may be considered permissive, since depending on the respective values of the PALT parameter for the control (PALT = 0.2) and treatment conditions, we can theoretically expect 20% (PALT = 0.4) and 40% (PALT = 0.6) changes in isoform proportions for the true DS genes. However, since FluxSimulator attempts to emulate actual sequencing experiments, the reads are generated after accounting for the commonly encountered technical biases during sequencing, which result in non-uniform sampling of reads along the lengths of the transcripts[23]. Therefore, as with real sequencing data, we do not expect perfect concordance between the original transcript-level counts and the observed event-level counts and hence allow for some deviation from the designated ΔPSIs by using a threshold of 10% to select the DS events.

### 2.3 ASTA-P Demonstrates a Good Trade-off Between Specificity and Sensitivity Compared with Other Methods in Computationally Simulated RNA-Seq Experiments

Salmon + DRIMseq and JUM appeared to be the superior methods as per the raw ROC metrics (**Figure 3, Table 1**). ASTA-P consistently showed the third best performance, following Salmon + DRIMseq and JUM.

**Figure 3.**
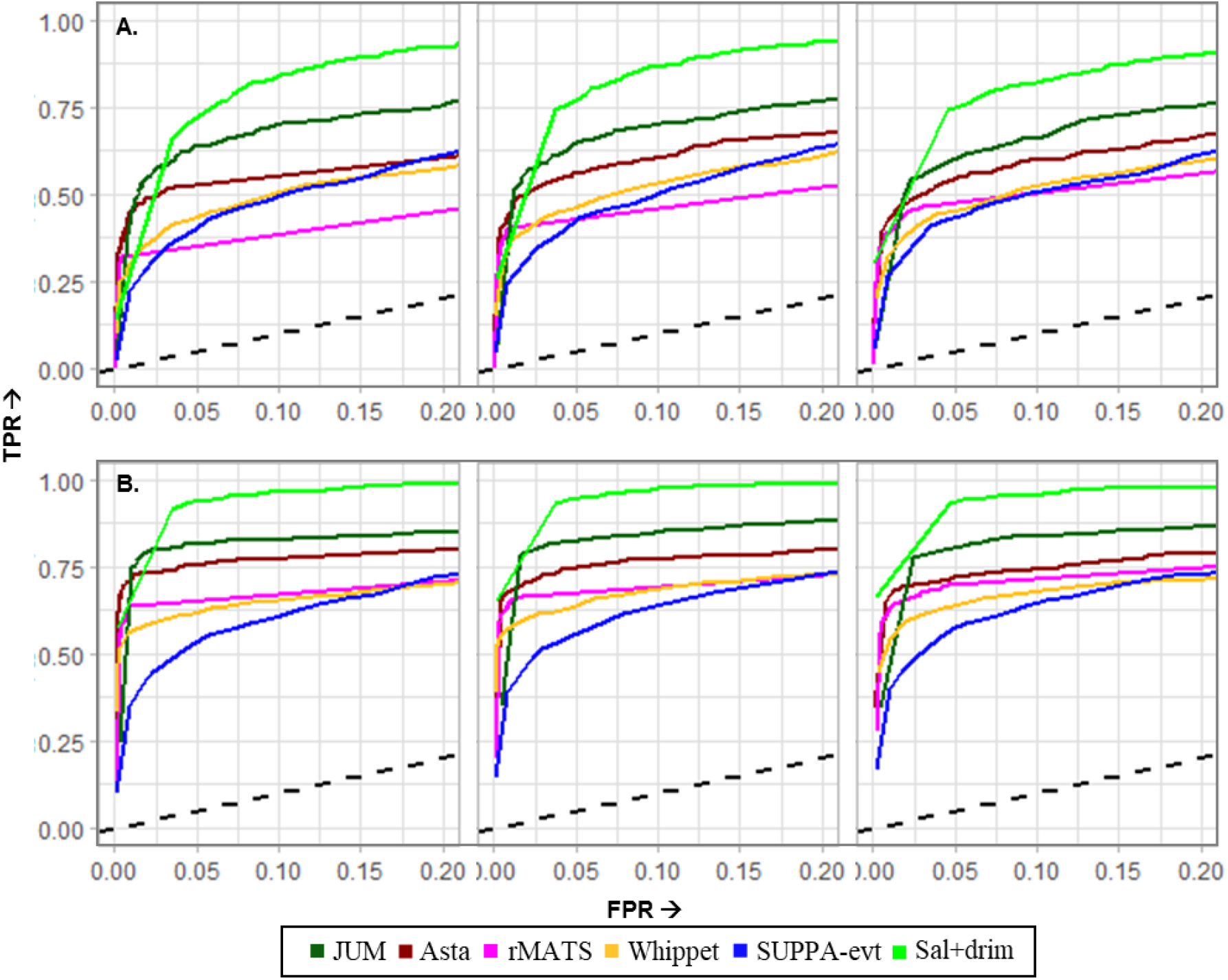
ROC curves evaluating the performance of the DS detection tools on the simulated DS genes with internal AS events. (A) PALT_treatment = 0.4 and (B) PALT_treatment = 0.6. In either panel, from left to right is increasing order of sequencing depth (15M, 30M, 60M).

The FDR-recall curves told a different story illustrating that ASTA-P and rMATS show the best FDR control, whereas Salmon + DRIMseq and JUM tend to sacrifice precision for sensitivity as both showed large positive deviations from the target FDR across all comparisons (**Figure 4**). Whippet and SUPPA also showed poor FDR control.

**Figure 4.**
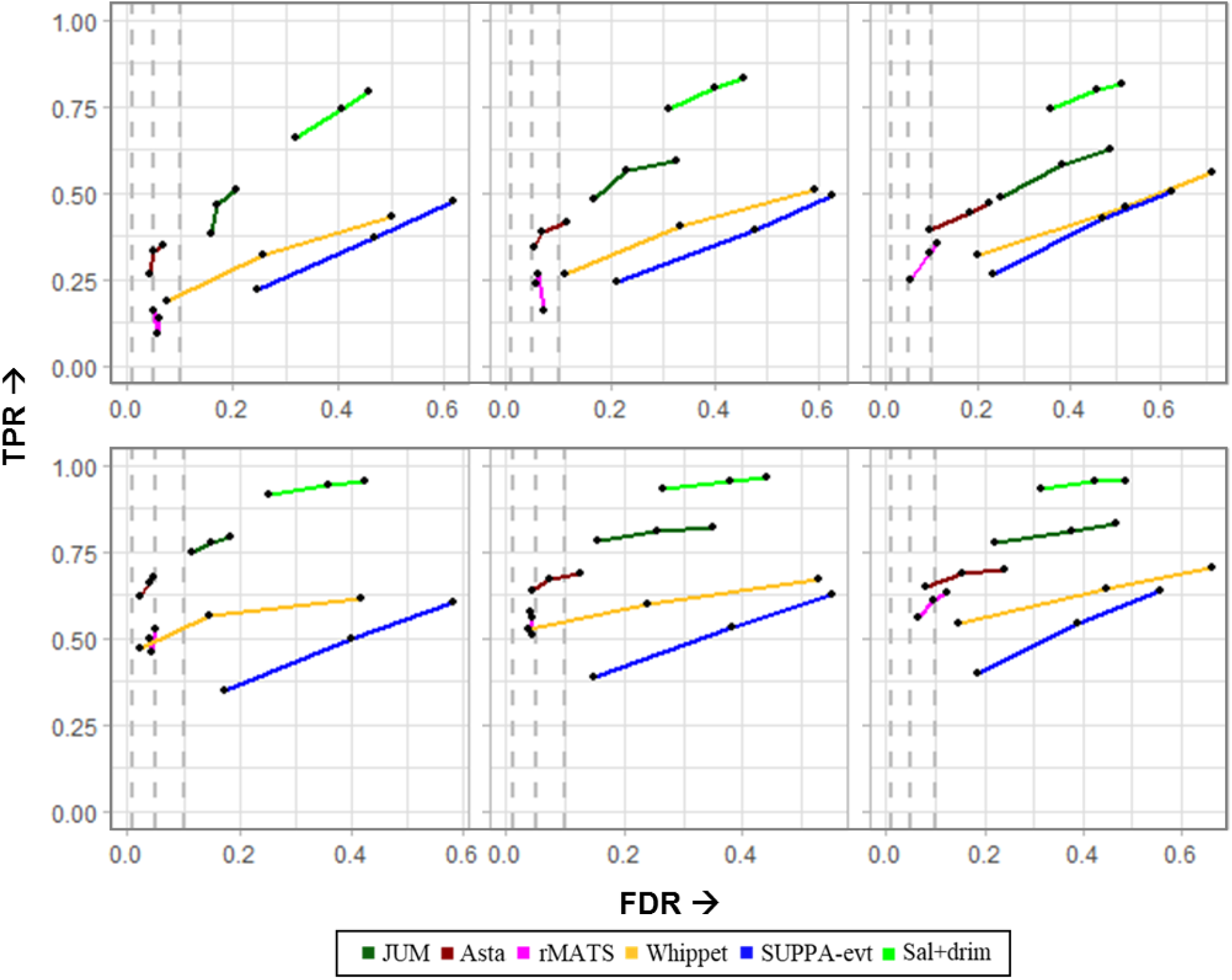
FDR-Recall curves evaluating the performance of the DS detection tools on the simulated DS genes with internal AS events at (A) PALT_treatment = 0.4 and (B) PALT_treatment = 0.6. In either panel, from left to right is increasing order of sequencing depth. The dashed grey lines represent the target FDR = {0.01, 0.05, 0.1}.

The discriminating power of all the tools improved considerably at the higher degree of DS (**Figure 3, Lower Panel**). Upon using the ΔPSI threshold for predicting DS genes, Whippet showed the strongest improvement in AUC for all comparisons (**Figure 5, Table 1**). It outperformed ASTA-P when PALT = 0.4, but their performances were comparable for PALT = 0.6. JUM and Salmon + DRIMseq still showed the best AUC, even though Salmon + DRIMseq’s performance dropped for the PALT = 0.4 comparisons. The AUCs for ASTA-P and Whippet approached that of JUM when PALT = 0.6 at the 60M sequencing depth. The FDR control for all tools also improved but ASTA-P and rMATS outperformed others as before.

**Figure 5.**
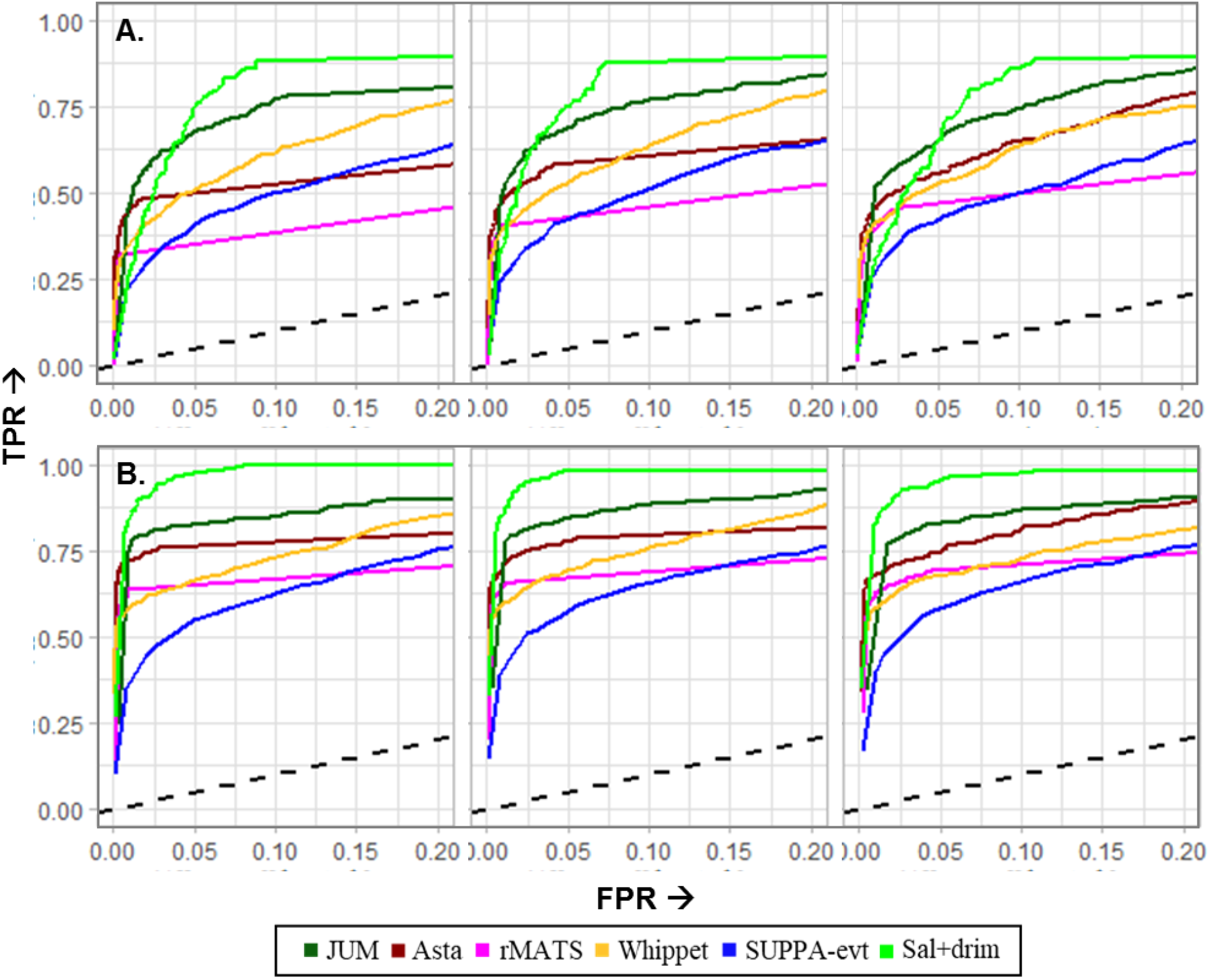
ROC curves evaluating the performance of the DS detection tools when applying the ΔPSI threshold to select predicted DS genes. (A) PALT_treatment = 0.4 and (B) PALT_treatment = 0.6. In either panel, from left to right is increasing order of sequencing depth (15M, 30M, and 60 M).

Sequencing depth exerted a variable effect depending on the tool and simulated degree of DS. ASTA-P showed and improvement with depth for all comparisons except when PALT = 0.6 without ΔPSI filtering. Only rMATS showed a consistent improvement in AUC with increasing depth across all comparisons (with and without ΔPSI thresholds). Sequencing depth did not appear to exert a beneficial effect on the AUC for the remaining tools. In terms of FDR control, the performance of all tools was negatively impacted with increasing depth (**Figure 6**).

**Figure 6.**
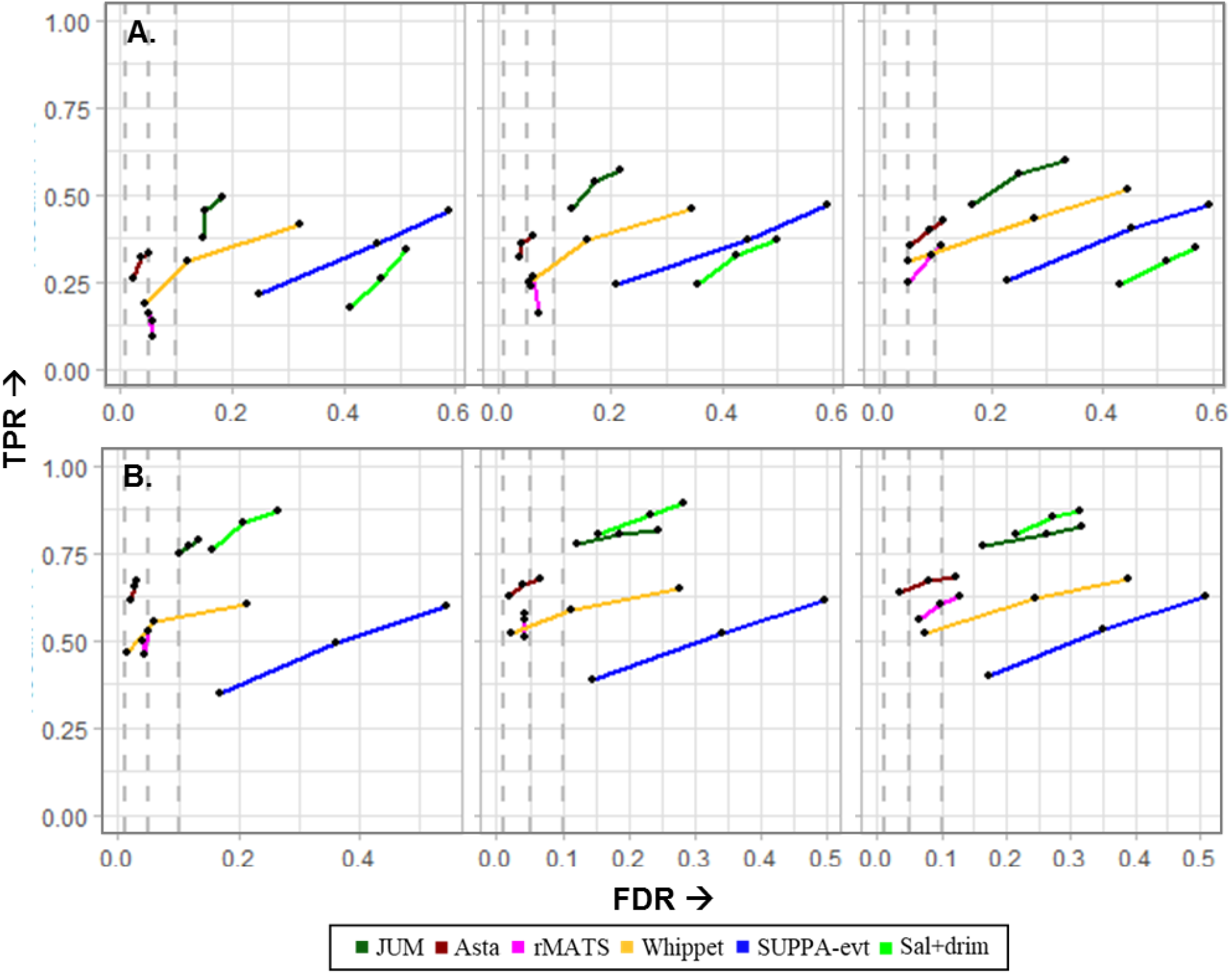
FDR-Recall curves evaluating the performance of the DS detection tools on the simulated DS genes with internal AS events, after applying the ΔPSI filter, at (A) PALT_treatment = 0.4 and (B) PALT_treatment = 0.6. In either panel, from left to right is increasing order of sequencing depth. The dashed grey lines represent the FDR = {0.01, 0.05, 0.1}.

### 2.4 Performance with an Incomplete Annotation

Even though >95% of human genes are estimated to undergo alternative splicing [1], the full repertoire of spliced isoforms has not been elucidated, and comprehensive RNA-seq analyses always report novel splice junctions. Additionally, for analyses involving disease samples, such as cancer samples, we can expect the expression of aberrant, novel splice forms. DS analysis tools must be able to deal effectively with such situations of “incomplete annotation”. Thus, in order to simulate this effect, we deliberately removed the chosen DS transcripts from the hg19 annotation and re-assessed the tools’ performances on the 60M depth data. ASTA-P (as well as JUM and Whippet) was able to maintain its AUC comparable to the corresponding value with the complete annotation (**Figure 7, Table 2**).

**Figure 7.**
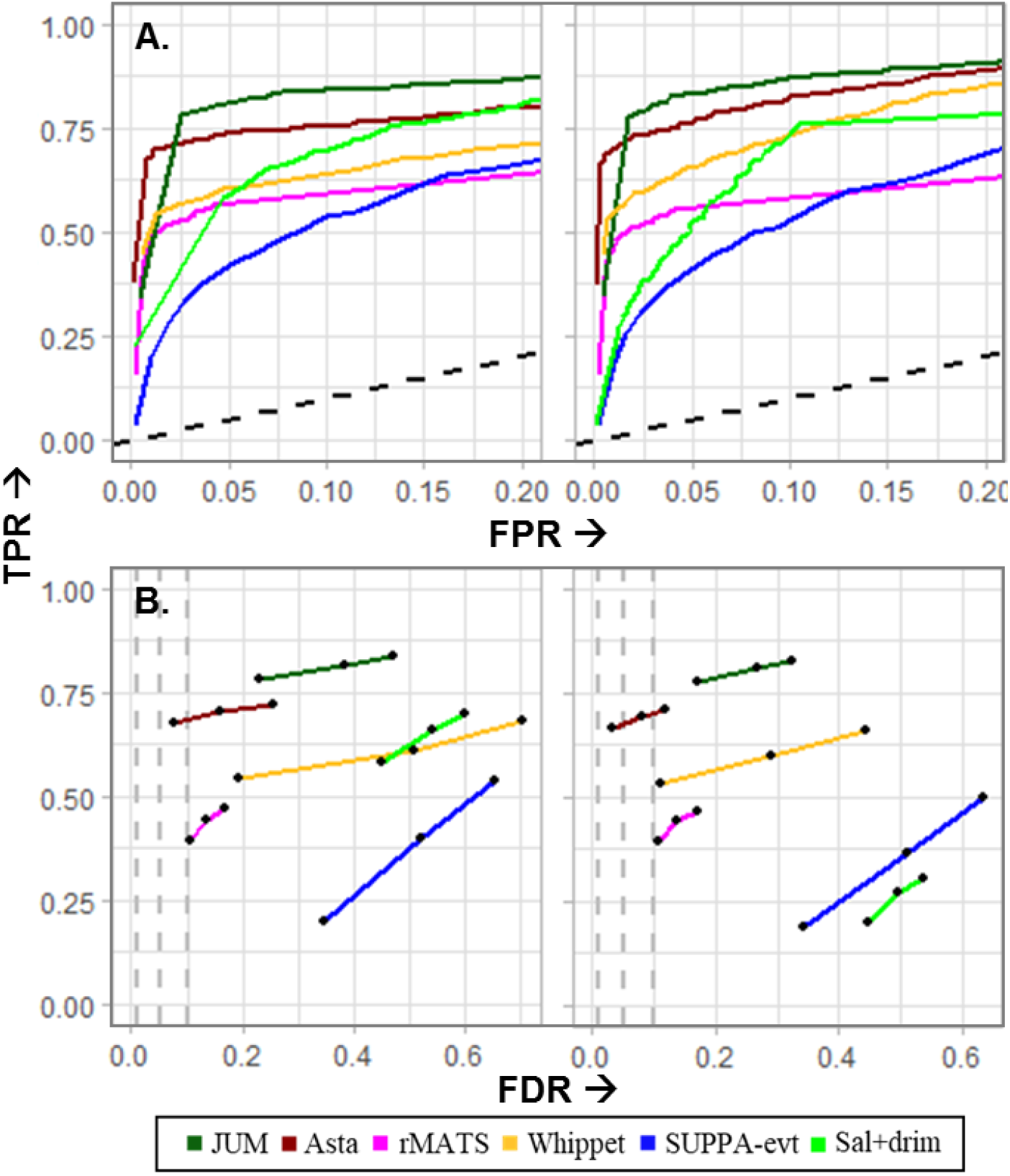
ROC curves describing the performance of the DS detection tools, for the 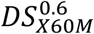 comparison, on (A) the full set of simulated DS genes and (B) on genes with internal AS events. In either panel, the left figures show the raw metric and the right figure shows the delta metric.

JUM had the best AUC with ASTA-P following in second. However, there was very little difference in the performance of the two after applying the ΔPSI filter. As expected, the performance of Salmon + DRIMseq worsened with an incomplete annotation across all comparisons. Interestingly, the loss in performance for SUPPA, which is also completely annotation dependent, was very modest. Even though rMATS has the functionality to discover novel events, its AUC also worsened. Concerning their FDR control, ASTA-P and rMATS performed best, but, as such, no method was able to achieve the target FDR. ASTA-P’s deviations were the lowest after applying the ΔPSI threshold.

### 2.5 Alternative Splicing Diversity in Cranial Neural Crest Cells

As mentioned in the introduction, we used ASTA-P to investigate arbitrarily complex AS events in delaminating human cranial neural crest cells (CNCCs). These cells arise in the neural tube during neurulation during nervous system development in vertebrates [25], [26]. Their subsequent delamination from the neural tube is critical for normal embryonic development. After delamination, bona fide migratory CNCCs arise and travel to different regions of the embryo to contribute towards the development of a wide array of bodily structures [27], [28]. The investigation of alternative splicing during NCC development has received very little attention, brought into the limelight through the study of CHARGE syndromes wherein congenital spliceosome mutations seem to impair the development of the CNCC derived tissues selectively [29]–[31]. Only very recently, independent studies examining the effects of deletion of splicing factors, specifically RBFOX2 [32] and ESRP1 [33], in mice embryos also reported the development of craniofacial defects due to aberrant splicing. During delamination, upregulation of the expression of the transcription factor, SOX10, establishes the identity of *bona fide* migratory CNCCs[34], [35]. Anu et al. constructed a SOX10:mMaple reporter hiPSC cell line and cultured them into CNCCs using their newly developed protocol (unpublished). The SOX10:mMaple reporter was used to select for pure populations of SOX10^+^ migratory NCCs as they appeared in culture in order to capture the delamination process. We used bulk RNA-seq data from sorted SOX10^−^ and SOX10^+^ cells to analyse the alternative and differential splicing patterns during this event. **Table S2** provides details of the RNA-seq samples.

Using STAR [24], Stringtie [36], and Salmon [13] in the annotation supplementation step of ASTA-P, we detected 64,536 transcripts, retained after filtering those expressed at a TPM >= 0.1 in at least five samples (smallest group size). We mined 23,997 splice events from these transcripts, including 17,053 (71%) classical 2D and 6944 (29%) non-classical type events (**Table S3**). After quantification and filtering of the event isoforms, we retained 19,353 “expressed events” including 15,285 classical 2D (79%) and 4068 (21%) non-classical type events (**Table 3**). Isoforms were filtered to retain those whose features (either unique or all) were supported by at least five reads in any five samples. The 19,353 events were subdivided into 14,844 Not-IR and 4509 IR events. We evaluated the different types of splicing patterns observed in these sets. Within the Not-IR events we detected 363 unique patterns, however, the classical 2D patterns were the most frequently used, accounting for 83.5% of the events (**Table 4A**). The diversity in splicing patterns was higher for IR events with 432 unique patterns, whereof 2D intron retention accounted for only 64% of the events (**Table 4B**). **Figure 8** depicts the top 20 most frequently observed non-classical splicing patterns in each category. Further, to gauge the effect of supplementing the reference transcriptome, we counted the number of events generating at least one isoform containing at least one unannotated junction. Splice junctions reported by STAR[24] were divided into five classes – annotated junctions (class 1), un-annotated junctions with annotated 5’ and 3’ splices sites (class 2), junctions with an un-annotated 5’ splice site (class 3), junctions with an un-annotated 3’ splice site (class 4), and un-annotated junctions where both 5’ and 3’ splice sites are un-annotated (class 5). We detected 5375 unannotated junctions (classes 2 - 5) supported by at least five uniquely mapped reads in any two samples (**Table S4**). Within the original set of 23,997 events, 1385 2D events (7.3%) and 747 HD events (14.2%) respectively contained at least one such unannotated junction (Table S4B). In the “expressed events” set, 1269 2D events (6.9%) and 296 HD events (24.6%) contained at least one un-annotated junction (**Table S5**). **Figure 8C** illustrates one such non-classical event in the coding region of *PRSS56* gene, which encodes a serine protease. The 5D event hosts four un-annotated junctions (coloured), two arise from inclusion of un-annotated 3’ acceptor sites (class 4), an un-annotated 5’ donor site (class 3), whereas one is an un-annotated combination of annotated splice-sites (class 2). This gene was recently shown to be involved in the generation of glial cells, which are one of the progenies of CNCCs, in mice [37].

**Figure 8.**
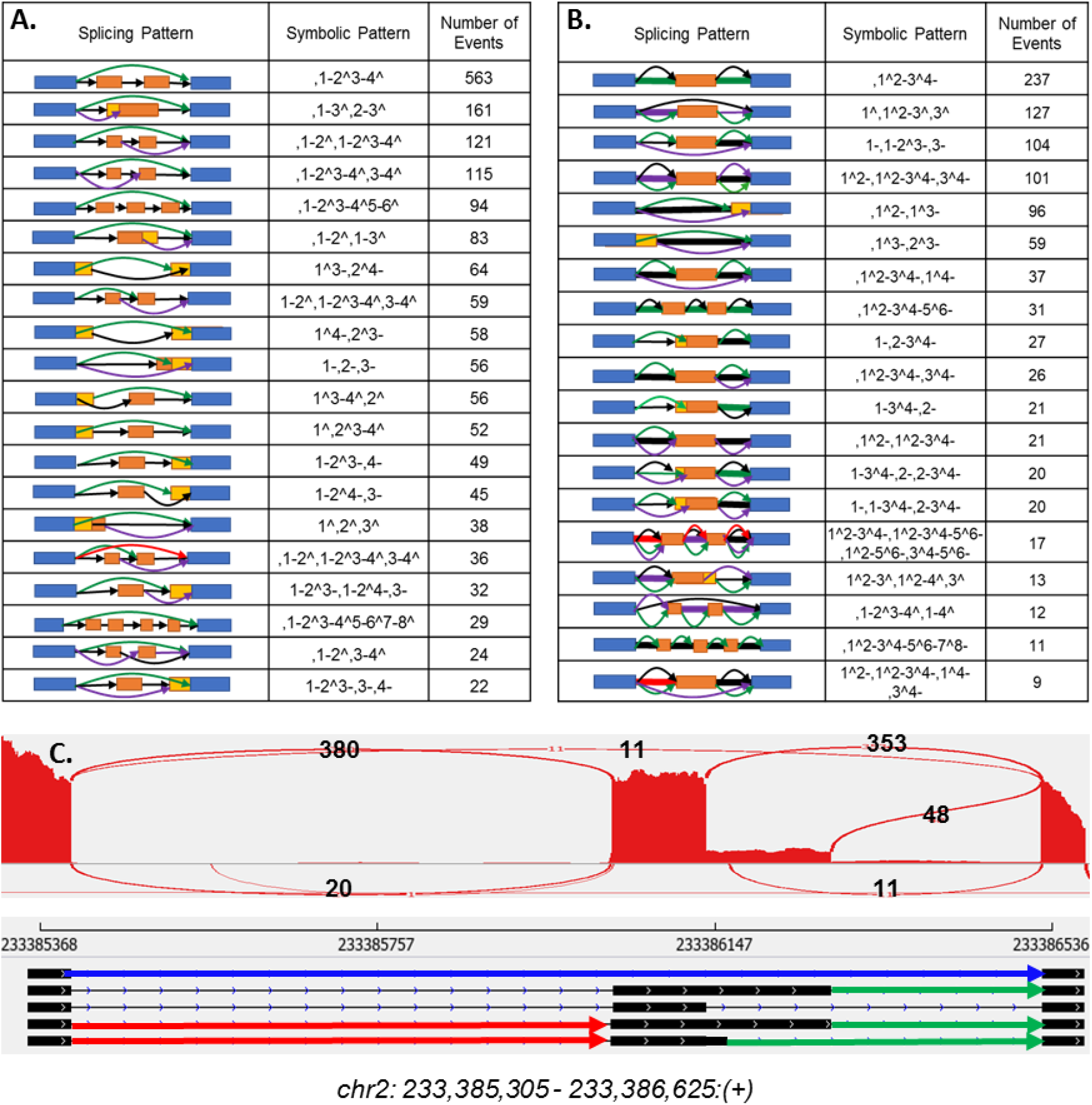
The top 20 most frequently detected non-classical splicing patterns detected in the (A) Not-IR and (B) IR events. (C) An example complex 5D event in the *PRSS56* gene, hosting multiple un-annotated junctions. (• Class 2, • Class 3, • Class 4 type un-annotated junction).

### 2.6 Isoform Usage and Noisy Splicing

Using DRIMSeq, we fit proportions for the Not-IR and IR events supported by total count of at least 20 reads in any five samples of at least one condition. 4994 (3373 Not-IR and 1621 IR; **Table 5**) events expressed at-least one alternatively spliced isoform, i.e. isoforms expressed at a proportion ϵ [10%, 90%] in at least one condition. This set comprised of 2142 non-classical events (174 2D-non-classical and 1968 HD). While the pervasiveness of AS across the human genome has been established, there is an ongoing debate whether rare splicing patterns and high dimensional splicing are functionally relevant (beyond the dominant isoforms) or merely observed due to stochastic mis-splicing producing low expressed transcripts[2], [38]–[41]. The “noisy-splicing” model argues that due to the extensive number of steps involved in a splicing reaction, occasionally, spurious transcripts are produced that manage to evade transcriptional quality control mechanisms, such as NMD[41]. It further proposes that the basal rate of generating these noisy transcripts depends on the fitness cost of the splicing errors and would be low for highly constrained genes, i.e. highly expressed (functionally important genes in a condition), and highly constrained regions within genes, i.e. the coding sequence of a gene[41], [42]. In order to explore these hypotheses in the context of our dataset, we first compared the total counts supporting the classical vs non-classical 2D patterns as well as the 2D vs HD events (**Figure 9A, B**). We did not detect any significant difference (p-value = 0.307) between the classical and non-classical 2D events, indicating that such splicing patterns are not restricted to low expressed genomic regions. For the HD vs 2D comparisons, HD events (d >= 3) were supported by significantly higher counts (p-value < 2.2e-16, for all tests) as compared to the 2D event counts. However, this observation may be expected since the detection of multiple isoforms would be more likely for highly expressed genes as compared to genes with lower expression levels.

**Figure 9.**
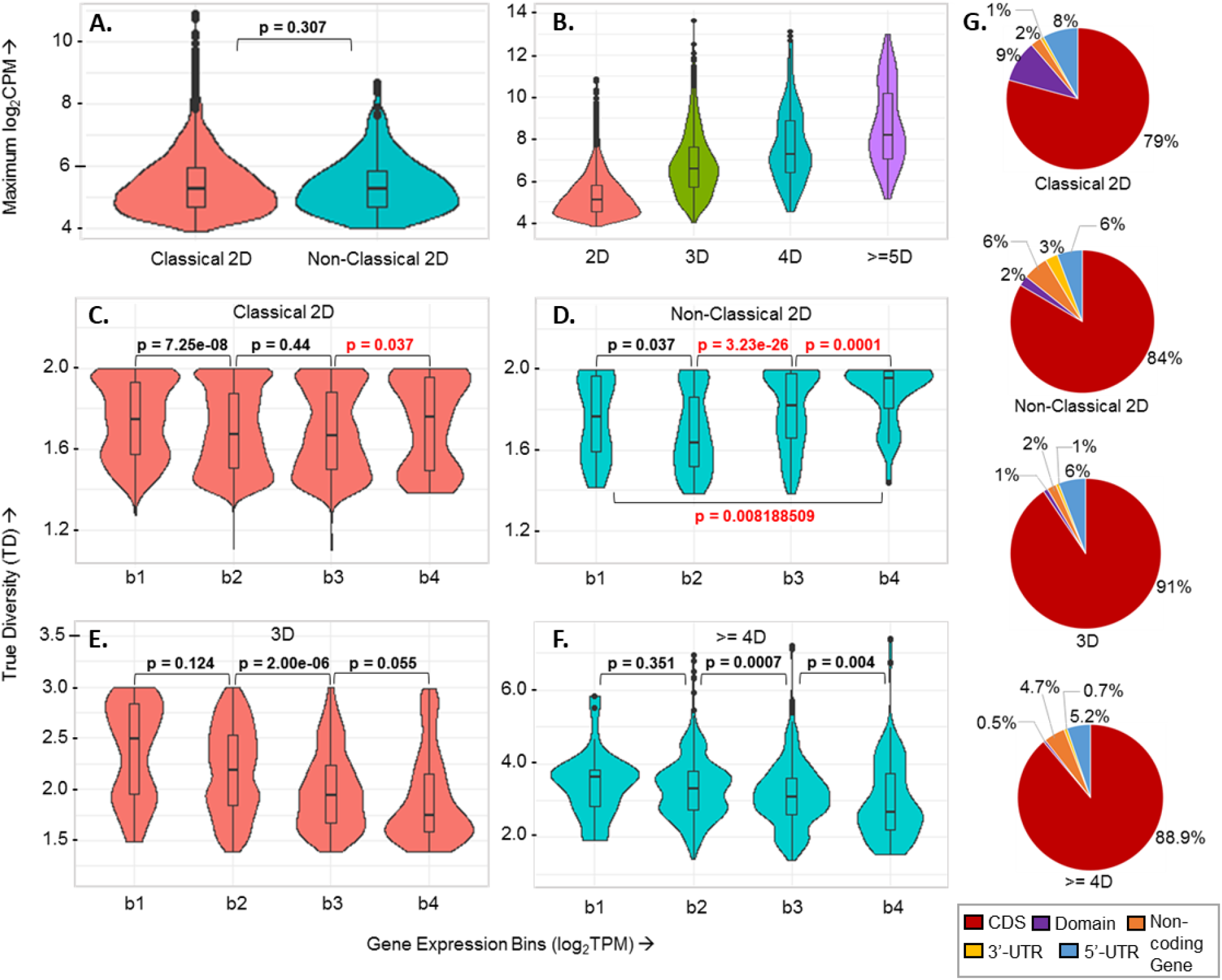
**(A)** Mean log2(CPM) total counts supporting 2D classical vs non-classical events. The mean is calculated across all replicates at any time point. The p-value represents the Wilcox test p-value from comparing the two sets of counts. **(B)** Mean log2(CPM) total counts supporting events of different dimensions. In this case, all consecutive pairwise comparisons using the Wilcox test had a p-value < 2.2e-16. **(C-F)** The True Diversity distributions of Classical and Non-Classical events grouped according to their host gene expression levels. P-values are from Wilcox test comparisons of TD values of events in consecutive bins. **(G)** Pie charts showing percent of Classical or Non-classical events lying in different gene regions (CDS, annotated domain within CDS, UTRs) or types (Coding vs Non-coding gene). **Gene Expression Bins: b1**: log2-TPM < 3, **b2**: 3 <= log2-TPM < 5, **b3**: 5 <= log2-TPM < 7, **b4**: log2-TPM >= 7

Therefore, next we analysed the isoform usage patterns for the HD (including non-classical 2D) events as compared to the classical 2D events. Usually, the Shannon entropy is a useful measure for describing the information content of a categorical probability distribution – i.e. a high entropy indicates a diffused distribution with multiple equiprobable states, whereas a low entropy is characteristic of a tight distribution with one highly frequent state [43]. However, when considering distributions with greater than 2 possible outcomes, such as the Dirichlet-multinomial in our case, the entropy measure is difficult to interpret owing to multiple probability states corresponding to very similar entropies and does not allow for a direct comparison of different dimensional distributions. Thus, we used the true diversity measure frequently employed in evolutionary biology studies, which estimates the number of equiprobable states that must exist in a system, in order to yield its corresponding entropy value [43]. For a d-dimensional event, the Shannon entropy (H) and True diversity (TD) are calculated as follows:

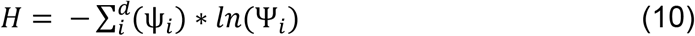

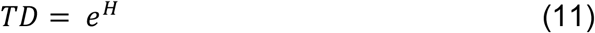

Here, ψ_i_ represents the proportion of any isoform generated from the event. Since the “noisy-splicing” model posits that the degree of AS would be lower for highly expressed genes, we can expect true diversities for the events in such genes to be lower than those for moderate or low expression gene events. We therefore binned the AS events into four bins according to the log transformed expression levels of their host genes and compared the true diversities of the events in each bin. As seen in **Figures 9C-F**, we observed variable trends with dimension. The 3D and >= 4D events agreed most strongly with the noise model – i.e. diversities significantly decreased with increasing gene expression. On the other hand, neither class of 2D events conformed to this trend. In fact, we observed the opposite trend for the non-classical 2D events with high diversity values becoming significantly more frequent for the most highly expressed genes, whereas for the classical 2D events, TD distributions became more pronouncedly bimodal with increasing gene expression. A GO analysis of 177 protein coding genes in the high expression bins (TPM >= 32; bins ϵ {b3, b4}), hosting 2D events with high TD values (>= 1.9) revealed an enrichment for a host of biological processes including RNA splicing, gene expression, cellular metabolic process, cellular biosynthetic process, translation and regulation of chromatin organization (**Tables 6, 7**, FDR < 0.05). Several genes (ATF4 [44], [45], CTNND1 [46], CTNNB1 [47], [48], HMGA1 [49], SMAD2 [50]) with established roles in regulating epithelial to mesenchymal transition (EMT), the process responsible for CNCC delamination, were included in this group, which contradicts the proposition of the noisy splicing model that potentially relevant genes should exhibit low degrees of alternative splicing. Further, a major fraction (85% = 175/206) of these high TD events, localized to CDS regions within their host genes and of those 126 (72% = 126/175) generated isoforms that were both supported by annotated protein coding transcripts. The second contention of the “noisy splicing” model is that AS events should more frequently be localized outside of coding regions. However, we did not find this to be the case, as at least 80% of the events across all dimensions, were found to lie in the CDS and annotated domain encoding regions (**Figure 9G**).

### 2.7 Differential Splicing During Neural Crest Delamination

In order to identify the differentially spliced events, we used DRIMSeq to compare the models: M_h_: Ψ ∼ Group ϵ {SOX10^+^, SOX10^−^} vs M_a_: Ψ ∼ Intercept, using the LRT. Only events expressed with a read count of 20 in at least five samples in both conditions were considered. DRIMSeq fits a multinomial model per gene (in our case event), as well as a beta-binomial model for each transcript (event-isoform) and provides independently FDR adjusted p-values at the event and isoform levels upon differential testing [11]. This is useful for determining both events and their specific differentially spliced isoforms. Since multiple sub-hypotheses are being tested per event, a stage-wise testing procedure, which aggregates evidence from all sub-hypotheses, allows for better overall FDR control, as opposed to independent adjustment of the standard event and isoform-level p-values[51]. Therefore, in order to determine the significantly differentially regulated isoforms in our data, we used the stageR package[51], which implements a two-stage testing procedure for performing differential transcript-level usage tests. This method involves a screening stage wherein events are filtered using the event-level adjusted p-value from DRIMSeq, followed by a confirmation stage where the significance threshold is re-adjusted according to the number of events that passed the screening stage. Multiple testing correction of isoform-level p-values is then performed across all hypotheses for an event to control the family wise error rate on this threshold. Using an overall FDR (OFDR) cut-off of 0.05, we identified 201 events (153 Not-IR, 48 IR events; **Table 8**) with significant changes (|ΔΨ| ≥ 10% for at least one isoform; FDR < 0.05). Of these, a significant fraction, 35% (n = 70/201) were non-classical events. Strikingly, the largest fraction of these events (49%) involved exons encoding parts of annotated protein domains within the host genes (**Figure 10**). A GO enrichment revealed that genes hosting these DS events are enriched for functions such as cytoskeletal organization, actin-filament based process, cell-cell junction organization, and cell migration, which are important for EMT (**Figure 10B, Table 9**). We defer from going into a detailed discussion of specific genes and events as this will be presented in a different publication.

**Figure 10.**
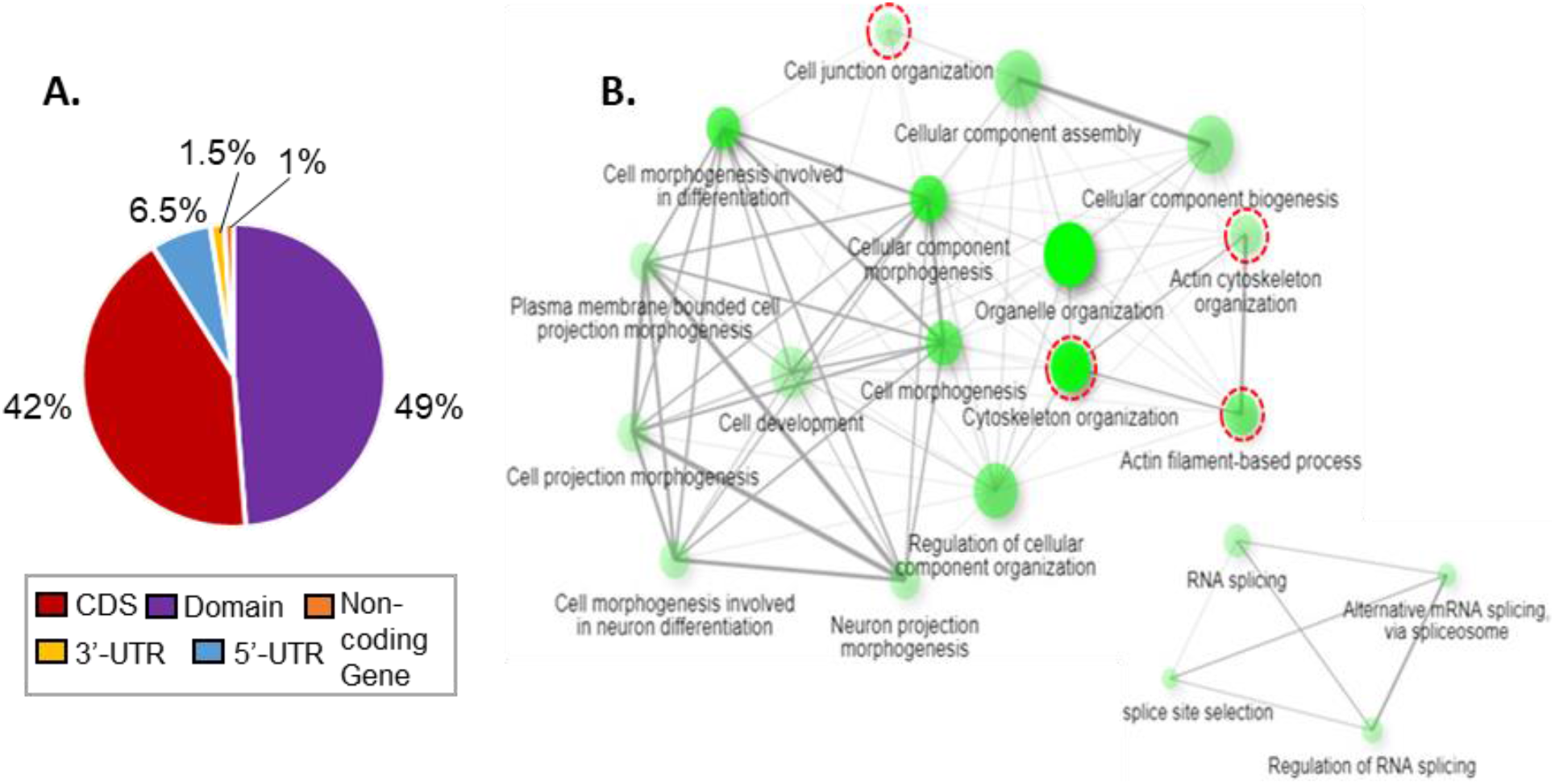
**(A)** The distribution of differentially spliced events in different regions of their host genes. **(B)** Network of a subset of top 20 Gene Ontology terms significantly enriched among the differentially spliced protein-coding genes. The encircled terms (red) are relevant for EMT.

### 2.8 The High Dimensional Splicing Code

The examination of sequence features of alternative exons/introns has helped in the identification of a “**Splicing Code**” [52]–[54] that can be used to predict splicing patterns for these events. We sought to identify whether any such features might be uniquely associated with non-classical 2D/HD events. In order to identify such features, we binned the AS events into five bins by dimension and examined features including the alternative exon lengths, the event outermost intron lengths, the event internal intron lengths, and the alternative exons 5’ and 3’ splice sites strengths. Apart from comparisons of the HD vs 2D exons, we also compared these against a control set of constitutively highly included exons (psi >= 95%) in both SOX10^−^ and SOX10^+^ samples.

Figure 11. shows the different features examined. We compared each feature using a Kruskal-Wallis test [55] (log_10_Feature ∼ Event Type ϵ {Constitutive, 2D-classical, 2D-non-classical, 3D, 4D, >= 5D}), followed by a Dunn post-hoc test [56]. Comparisons of alternative exons and outermost flanking introns lengths between event types did not reveal any meaningful patterns. However, the internal introns lengths appeared significantly increased with dimension among the alternative events. In humans, shorter exons with weak splice sites and flanked by longer introns are spliced less efficiently [57]–[59]. However, comparing against the constitutive exons, only the >= 5D events contained significantly longer internal introns, whereas introns of both classes of 2D events were in fact significantly shorter. With respect to the splice-sites, alternative events across all dimensions except 2D-non-classical harboured significantly weaker 3’ splice sites as compared to constitutive events. A similar, but weaker trend was apparent for the 5’ splice sites as well.

**Figure 11.**
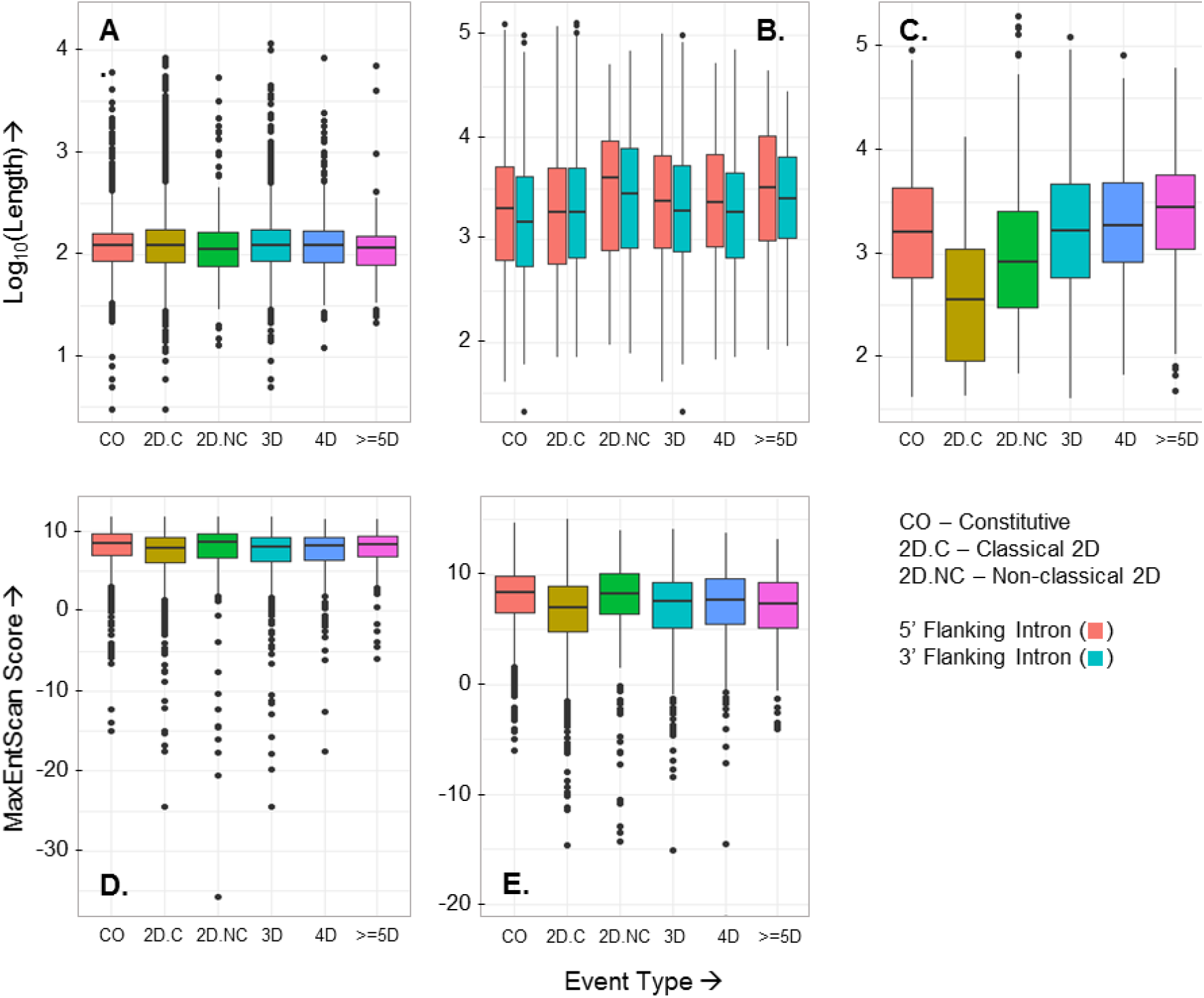
The different sequence features examined for determining an HD splicing code: **(A)** Log-transformed alternative exon lengths. **(B)** Log-transformed lengths of the events 5’ and 3’ flanking introns. **(C)** Log-transformed lengths of events internal introns. **(D-E)** Maximum entropy scores of the 5’ donor and 3’ acceptor sites of alternative exons.

### 2.9 Discussion

Since the first evidence of AS emerged around four decades ago, it is now estimated that around 95% of human genes undergo alternative splicing to produce multiple mRNA isoforms [1]. The advancement in experimental techniques to comprehensively survey the transcriptome was accompanied by the development of computational methods to detect and quantify these splice forms from the transcriptomic data. Among these methods, full-length transcript level analyses are clearly the objective but are presently limited in accuracy due to experimental limitations. A greater focus therefore has been on the development of event level methods, which quantify a set of pre-defined binary splicing patterns using uniquely assignable junction specific RNA-seq reads allowing for a more accurate analysis[16], [60]. In this paper, we described such an event-based workflow, ASTA-P. Rather than limiting ourselves to the annotated transcriptome and classical 2D splicing patterns, we employed a hybrid approach combining full-length transcript reconstruction to enrich the annotation, followed by splicing graph assembly, which is then traversed to mine arbitrarily complex splicing variations. Using a simulated dataset, we compared our pipeline to four popular events-based (Whippet, JUM, rMATS, and SUPPA) and one transcript-based (Salmon +DRIMseq) splicing analysis methods w.r.t. their abilities to detected differentially spliced events, examining the effects of differential splicing degree, sequencing depth, and incomplete annotation. We also examined how the usage of a ΔPSI threshold, often used to select DS genes in splicing analyses, impacts the performance of these tools. Transcript-based methods are not considered the most suitable option for DS analysis because of quantification inaccuracies that may arise due to the inherent limitations of short read RNA-seq data. This was somewhat reflected in the performance of the Salmon + DRIMseq method as it showed a high recall but low precision. Further, in samples with the lower simulated degree of splicing change (PALT = 0.4), its recall dropped significantly upon use of the ΔPSI threshold suggesting that it is able to capture a change in splicing proportions but the ΔPSI value may not be captured effectively for smaller true changes in splicing. The recently published events-based method, JUM, showed a similar performance to Salmon + DRIMseq, i.e. high recall but low precision. However, like other events-based tools, its recall and FDR control improved upon application of the ΔPSI threshold regardless of the simulated degree of DS, indicating that it must be used with careful parameter calibration to reduce the FDR. Our pipeline, ASTA-P, was most frequently the third best in terms of recall and along with rMATS was able to achieve the best FDR control. Whippet generally followed in after ASTA-P in terms of recall but had poor FDR control. SUPPA consistently showed the worst FDR control. Whereas the degree of differential splicing had a clear effect on all tools’ performance, i.e. both recall and FDR control improved with greater simulated DS degree, sequencing depth did not have an appreciable effect on recall for any tools except rMATS whose recall consistently improved with increasing depth. FDR control was impaired consistently for all tools with increasing depth. Lastly, ASTA-P, JUM and Whippet appeared to be robust to incomplete annotation, both in terms of recall and FDR control. rMATS’ dependence on annotation was uncovered in this analysis as it suffered a greater loss in recall even compared to the purely annotation dependent tool, SUPPA and showed reduced FDR control. Thus, the sensitivity of ASTA-P must be improved, but currently it offers a reasonable trade-off between discovery and accuracy. Further, we applied ASTA-P to analyse alternative splicing patters in bulk RNA-seq data from hiPSC generated cranial neural crest cells (CNCCs), simulating the embryonic event of CNCC delamination from the neural tube in order to migrate and give rise to different body structures [26]. We examined the types and properties of splicing events detected in this data. We found a remarkable diversity of 795 unique splicing patters across 19,353 events. Among these, non-classical 2D patterns and HD splicing accounted for 21 % of the events detected. Such events also comprised a significant fraction (43%) of the 4994 alternatively spliced events, expressing at least one isoform at a proportion ϵ [10%, 90%] in any condition. A much-debated question is multiple spliced isoforms of a gene actually contribute towards expanding the transcriptome and proteome [2]. Noisy splicing models contend that, for most genes, minor isoforms are stochastic junk and their expression depends on the resource cost of mis-splicing [41]. As a first step, we did not find non-classical 2D or HD isoforms to be supported by lower number of total reads as compared to the classical 2D patterns. Further, using the notion of “True Diversity”, we demonstrated that indeed for the HD splicing events, isoform usage does agree with the noisy splicing model getting more concentrated towards a single dominant isoform for highly expressed genes. Although, even for the most highly expressed genes, a major fraction of these events had a true diversity >= 2, suggestive of significant expression beyond the most dominant isoform. Interestingly, 2D events defied the propositions of the noisy splicing models, with high diversity values becoming more common with increasing gene expression. Furthermore, a subset of these AS events hosted by genes linked with EMT and other functions such as regulation of gene expression, mRNA splicing, cell-cycle regulation, co-expressed multiple major isoforms. Similar observations regarding isoform co-expression in genes encoding transcription factors were also made in a recent study examining alternative splicing across a vast set of tissues and cell lines[61]. The recently published high-complexity splicing analysis tool, Whippet also reported that around 40% of protein coding genes express multiple major isoforms [10]. Non-classical type events also contributed significantly towards regulation, with 35% of the regulated events (|ΔPSI| >= 10%, FDR < 0.05) falling into this category. These events were detected in genes participating in process, such as cytoskeletal organization, actin-filament based process, cell-cell junction organization, that are relevant for the EMT that occurs during CNCC delamination.

## 3. Methods

### 3.1 SIMULATED DATA ANALYSIS

#### 3.1.1 RNA-Seq Simulations

In order to evaluate the performance of the tools, we generated artificial RNA-seq data using FluxSimulator [23], according to a previously described workflow[22]. We based the simulated data on bulk RNA-seq data from hiPSC derived cranial neural crest cells expressing different levels of the transcription factor, SOX10. In order to perform a two-group comparison for the differential splicing evaluation, we modelled two groups of data, i.e. control and treatment based on the SOX10^−^ and SOX10^+^ neural crest datasets respectively. As per the workflow, gene-level counts for each condition were sampled from NB distributions estimated using the SOX10^−^ and SOX10^+^ datasets. These gene-level counts were compared with the supplied annotation (Ensembl.v75) to identify the expressed genes and a random subset of 2000 multi-transcript genes were selected as the “true DS genes” from those. For each of the true DS genes, a transcript was selected at random as the main DS transcript, and its proportion was set equal to a PALT parameter, defined for a given condition. The remaining proportion, (1 - PALT), was equally distributed among the remaining transcripts. Depending on the value of the PALT parameter chosen for the control and treatment conditions, we were able to simulate different degrees of DS. For each of the “true non-DS genes”, transcript proportions were sampled once, from a uniform distribution, and were fixed at the same values across all conditions. Further, a coverage parameter, c, was used to control the depth of the simulated RNA-seq data. The scripts and definitions of the associated parameters for running the simulation workflow are available at: https://github.com/ruolin/ASmethodsBenchmarking. The exact commands we used for running our simulations are provided in **Supplementary Section S1**.

#### 3.3.2 Differential splicing detection comparisons

The parameters used for running each tool are provided in **Supplementary Section S2**. We evaluated the performance of the tools in detecting differential splicing in terms of their discriminative power and ability to control the false discovery rate (FDR). For this purpose, we plotted the receiver operator characteristics (ROC) curves and FDR vs Recall curves[62], [63]. If TDS, TNDS respectively denote the set of true DS and true non-DS genes and, PDS and PNDS respectively denote the genes predicted to be significantly DS or undergoing no significant DS by a tool, the following set of metrics can be computed:

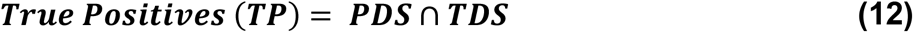

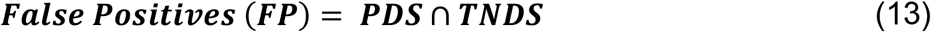

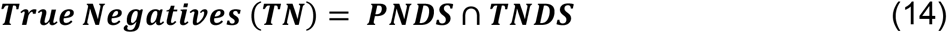

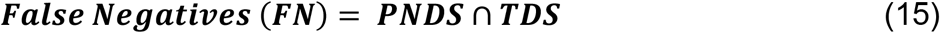

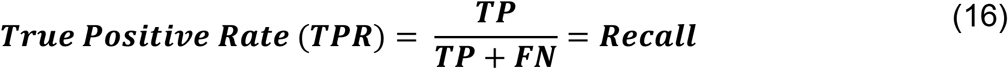

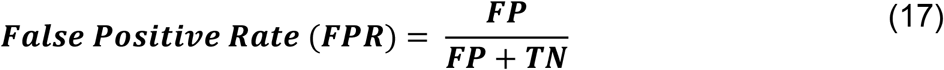

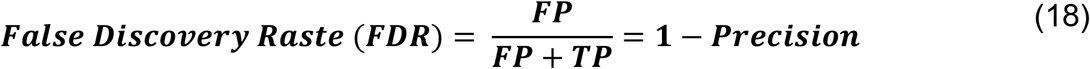

In order to compute the PDS and PNDS set, each gene is assigned a “score” and a threshold value (t) is used to classify each gene as DS (score >= t) or non-DS (score < t). ASTA-P, rMATS, JUM, SUPPA and Salmon + DRIMseq compute an adjusted p-value (p-adj) of differential splicing for each event. For these tools, for each gene, we selected the event with the lowest p-adj and assigned it a differential splicing probability score = (1 – p-adj). On the other hand, Whippet outputs a probability of differential splicing for each node, as opposed to a significance p-value, and we used this probability as it is. For each gene, the node with greatest probability of DS was selected for the evaluation. To plot the ROC (TPR-FPR) and FDR-Recall curves, the above metrics were computed across a range of DS probability score thresholds (ϵ [0, 1]). For the ROCs curves, the area under the curve (AUC) was further computed using the ROCR R-package [64].

### 3.2 REAL DATA ANALYSIS

#### 3.2.1 Pre-processing of RNA seq samples

Poly-A selected mRNA was sequenced from six biological replicate samples each of SOX10^−^ and SOX10^+^ NCCs to produce 100bp, stranded, paired-end reads. Unfortunately, one of the SOX10^+^ samples was compromised in quality and was discarded from further analysis. The FASTQ formatted read files were assured to be of high quality using FASTQC [65]. Reads were aligned to the hg19 genome build using the splice aware aligner STAR (v2.6.1) [24]. Since Salmon does not perform de-novo transcript reconstruction, Stringtie [66] was used to detect any novel transcripts expressed in our data. As recommended by the Stringtie authors in [66], we perform annotation guided reconstruction using the Ensembl (version 75) [67] annotation for each condition (SOX10^-^ and SOX10^+^) and then merge the individual transcriptomes to generate a non-redundant set of transcripts. Due to the low confidence in correctly detecting alternative transcription start and end sites, Stringtie merges any predicted transcript with a reference transcript if they share the same internal intron-exon structure [66]. We do not attempt to detect alternative transcription start sites (TSSs) or alternative poly-adenylation events in this data and focus only on alternative splicing events. Salmon was used to quantify the final merged transcriptome. Transcripts were filtered to keep only those expressed at a level of at least 0.1 TPM in at least five samples (smallest group size).

#### 3.2.2 Detection and Quantification of AS events

Detection and quantification of the AS events was performed according to the pipeline detailed in Section 2.2. For filtering the splicing events’ isoforms, at least five reads were required to uniquely map to a unique feature of an isoform in at least five samples. Isoforms without any unique features were retained if all their junctions were expressed with at least five reads in at least five samples.

#### 3.2.3 Differential Splicing Analysis

We filtered the events to retain those expressed with total read count of 20 in at least five samples of both conditions. We modelled the isoform counts for each event using DRIMSeq, and identified the differentially spliced events by comparing the full model (∼ Group ϵ [SOX10^−^, SOX10^+^]) to the null model (intercept only) using a likelihood ratio test. To further identify the differentially regulated isoforms of these events, we used the stageR package [51] to assign FDR values at the isoform level. Significantly differentially spliced isoforms were identified using an event level p-value threshold of 0.05, as those with a |ΔPSI| >= 10% and an isoform level FDR < 0.05.

#### 3.2.4 Feature Analysis of Alternatively Spliced Events

AS exons were identified selected according to their inclusion level, i.e. Ψ_exon_ ϵ [10%, 90%] in at least one condition. The inclusion level for an exon of any event was determined as the ration of the cumulative proportion of isoforms containing that exon vs. all isoforms generated through that event. We binned these exons by type and dimension of their host events. We compared their log-transformed lengths and the maximum entropy scores of their splice sites against each other as well as a set of constitutively included exons (Ψ_exon_ > 90% in both SOX10^-/+^ cells). We used the MaxEntScan tool [68] to score the splice sites. For comparing introns, for each event, we extracted the outermost 5’ and 3’ introns as flanking introns and compared their log-transformed lengths. We also extracted internal (non-flanking) introns to compare their log-transformed lengths between different event types.

#### 3.2.5 Functional Analyses of Differentially Spliced Events

In order to assign events to gene regions, we compared event coordinates to the coordinates of different functional regions (5’-UTR, CDS, and 3’-UTR) within the principal coding transcript for each gene, indicated in the APPRIS database [69]. For events lying in CDS regions, we further determined their overlap with annotated Pfam [70] and Uniprot [71] domain records available through the UCSC table browser [72] to assign them to “Domain” regions. For functional interpretation of the differentially spliced genes, we conducted a GO enrichment analysis using STRING browser.

## Supplementary Sections

### S1. Commands for RNA-seq Simulations

#### Modelling the real CNCC dataset, and selecting DS genes

~~~
python **cal_NB_counts.py** Ensembl.gff3 -g1 24707.sam 24708.sam -g2 24705.sam 24706.sam -m AS-genes
~~~

#### Running Flux Simulator

~~~
PARF=“Location/of/FluxSimulator”
   **python generate_rnaseq.py group1.nbcounts** AS_genes_list.txt
   $PARF”/”myPara.par g1.15M.0.2 **-p 0.2 -c 50
   python generate_rnaseq.py group2.nbcounts** AS_genes_list.txt
   $PARF”/”myPara.par g2.15M.0.4 **-p 0.4 -c 50
   myPara.par** is the FluxSimulator parameter file
   **p** indicates the proportion of the selected DS transcript in this dataset
   **c** indicates coverage, which in turn dictates depth. c = 50 here results in a depth of 15M reads
   **g1(2).15M.0.2(4)** are output prefixes for the generated sequencing files.
~~~

The scripts, calc_NB_counts.py and generate_rnaseq.py are available at https://github.com/ruolin/ASmethodsBenchmarking.

### S2. Tool-Wise Commands

#### ASTA-P

##### 1.1 Annotation Supplementation with STAR and Stringtie

~~~
For n in (1…3)
  **STAR** --runMode alignReads --runThreadN 20 --genomeDir $GEN \
  --genomeLoad NoSharedMemory --readFilesIn **G1.Sn.R1.fq.gz G1.Sn.R2.fq.gz** \
  --readFilesCommand gunzip –c \
  --outFileNamePrefix **G1.n** \
  --outSAMattributes NH HI NM MD AS nM jM jI XS \
  --outSAMtype BAM SortedByCoordinate \
  --limitBAMsortRAM 37125051484
  **STAR** --runMode alignReads --runThreadN 20 --genomeDir $GEN \
  --genomeLoad NoSharedMemory --readFilesIn **G2.Sn.R1.fq.gz G2.Sn.R2.fq.gz** \
  --readFilesCommand gunzip –c \
  --outFileNamePrefix **G2.n** \
  --outSAMattributes NH HI NM MD AS nM jM jI XS \
  --outSAMtype BAM SortedByCoordinate \
  --limitBAMsortRAM 37125051484
  GEN=“/Path/to/STAR.Index”
~~~

For n in (1…3)

~~~
**stringtie G1.nAligned.sortedByCoord.out.bam** -o **G1.n.f0.01.gtf** -p 20 -G
$REF -eB -f 0.01
**stringtie G2.nAligned.sortedByCoord.out.bam** -o **G2.n.f0.01.gtf** -p 20 -G
$REF -eB -f 0.01
REF=“/Path/to/EnsemblAnnotation.gtf”
**stringtie** --merge -f 0.01 -G $REF -p 20 -o G12.f0.01.gtf assembly_GTF_list.txt
assembly_GTF_list.txt contains paths to the sample-wise assemblies generated in the previous step.
~~~

##### 1.2 Transcript filtering, Event Mining, and Isoform Filtering

This step involves quantification of the merged set of transcripts (G12.f0.01.gtf) using Salmon with default parameters, followed by filtering to keep transcripts expressed at 0.1 TPM in at least 3 samples (smallest group size in the simulation study).

Example Salmon Command:

~~~
**gffread** -w G12.f0.01.fa -g $GEN G12.f0.01.gtf
**salmon** index -t G12.f0.01.fa -i $IDX -p 20
GEN=“/Path/to/genomefastafiles”
IDX=“/output/path/for/storing/index”
**salmon** quant -i $IDX -l A \
  -1 G1.S1.R1.fq.gz \
  -2 G1.S1.R2.fq.gz \
  -p 20 --validateMappings --rangeFactorizationBins 4 --gcBias --seqBias –o G1.1
~~~

Transcript expression tables are compiled and filtered using Tximport in R.

Filtered transcripts File: G12.f0.01.filt.gtf

**astalavista** -t asta -i G12.f0.01.filt.gtf -e [ASI] -d 0 -o G12.f0.01.filt.asta.gtf.gz

After this step, the full set of events (G12.f0.01.filt.asta.gtf.gz) are separated into Not-IR and IR events, quantified, and filtered using custom python scripts.

For the simulation analyses, event isoforms were filtered to keep those expressing at least one unique feature or all features with 5 (30M, 60M depth) or 2 (15M depth) reads.

Output Files:

Event Details: G12.NotIR.allD.final.e2g.out, G12.IR.allD.final.e2g.out Counts: G12.NotIR.final.IsoExprsn.out, G12.IR.final.IsoExprsn.out

##### 1.3 DRIMseq and StageR Commands

~~~
Samples tables is formatted as: “Sample Group”, where Group ϵ {G1, G2} txiT = G12.NotIR.final.IsoExprsn.out, G12.IR.final.IsoExprsn.out
drimObj <- **dmDSdata**(counts = txiT, samples = samples)
drimObj <- **dmFilter**(drimObj, min_samps_gene_expr = 3,
      min_gene_expr = 20)
model.full <- model.matrix(∼Group, data = samples)
drimObj <- **dmPrecision**(drimObj, design = model.full, verbose = 2, add_uniform = T)
drimObj <- **dmFit**(drimObj, design = model.full, verbose = 2, add_uniform = T)
   dtest.hcl <- **dmTest**(drimObj, coef = “GroupG2”)
   res.hcl <- results(dtest.hcl)
   res.hcl$adj_pvalue[is.na(res.hcl$adj_pvalue)] <- 1
   res.hcl.ft <- results(dtest.hcl, level = “feature”)
   pConfirmation <- matrix(res.hcl.ft$pvalue, ncol = 1)
   pScreen <- res.hcl$adj_pvalue
   sRObj.clcm <- **stageRTx**(pScreen = pScreen, pConfirmation = pConfirmation, pScreenAdjusted=TRUE, tx2gene=tx2gene)
   sRObj.clcm <- **stageWiseAdjustment**(object = sRObj.clcm, method=“dtu”, alpha=0.05)
~~~

#### rMATS

~~~
**python rmats.py** --b1 b1.txt --b2 b2.txt --gtf $GTF --od $OP -t paired --readLength 100 --cstat 0.05 --libType fr-secondstrand
$GTF=“path/to/ EnsemblAnnotation.gtf”
b1.txt contains paths to group 1 BAM files
b2.txt contains paths to group 2 BAM files
~~~

#### Whippet

##### 1. Building Whippet Index

~~~
**julia whippet-index.jl** --fasta $GEN --bam G12.sort.rmdup.bam --gtf $GTF
$GTF=“/path/to/EnsemblAnnotation.gtf”
$GEN=“/path/to/genomeFastaFiles”
G12.sort.rmdup.bam – As recommended by Whippet authors (https://github.com/timbitz/Whippet.jl) merged, sorted and duplicate filtered read alignments from all samples.
~~~

##### 2. Running Whippet Quant and PSI differential analysis

For n in (1…3)

~~~
 **julia whippet-quant.jl** G1.Sn.R1.fq.gz G1.Sn.R2.fq.gz -o G1.n
 **julia whippet-quant.jl** G2.Sn.R1.fq.gz G2.Sn.R2.fq.gz -o G2.n
**julia whippet-delta.jl** -a G1.1.psi.gz,G1.2.psi.gz,G1.3.psi.gz -b G2.1.psi.gz,G2.2.psi.gz,G2.3.psi.gz -o outputDir
~~~

**JUM: JUM contains stepwise scripts (containing commands) that are run in order (https://github.com/qqwang-berkeley/JUM/wiki/3.1.-Manual-running-JUM-(v2.0.2-and-up)). I ran those with default parameters for all datasets**.

#### SUPPA

##### 1. SUPPA extract splicing events

~~~
**python suppa.py generateEvents** -i $REF -o $OP/hg19_suppa_event_level
-f ioe -e SE SS MX RI FL
~~~

##### 2. SUPPA Quant and Differential analysis

for ev in {SE, A3, A5, MX, RI, AF, AL}

~~~
**python suppa.py psiPerEvent** --ioe-file
hg19_suppa_event_level_$ev_strict.ioe --expression-file G1.tpms.out -o
G1.$ev.out
**python suppa.py psiPerEvent** --ioe-file hg19_suppa_event_level_$ev_strict.ioe --expression-file G2.tpms.out -o
G2.$ev.out
**python suppa.py diffSplice** --method empirical --input
hg19_suppa_event_level_$ev_strict.ioe --psi G1.$ev.out.psi G2.$ev.out.psi\
--tpm G1.tpms.out G2.tpms.out --area 1000 --lower-bound 0.05 -gc -o
DSE.$ev.out
~~~

### Salmon

#### 1. Salmon Quant

~~~
**gffread** -w Ensembl.fa -g $GEN Emsembl.gtf
**salmon** index -t Ensembl.fa -i $IDX -p 20
GEN=“/Path/to/genomefastafiles”
IDX=“/output/path/for/storing/index”
~~~

For n in (1…3)

~~~
**salmon** quant -i $IDX -l A \
-1 G1.Sn.R1.fq.gz \
-2 G1.Sn.R2.fq.gz \
-p 20 --validateMappings --rangeFactorizationBins 4 --gcBias --seqBias –o
G1.n
**salmon** quant -i $IDX -l A \
-1 G2.Sn.R1.fq.gz \
-2 G2.Sn.R2.fq.gz \
-p 20 --validateMappings --rangeFactorizationBins 4 --gcBias --seqBias –o
G2.n
~~~

Transcript expression tables are compiled using Tximport in R.

#### 2. DRIM-seq Commands

Samples tables is formatted as: “Sample Group”, where Group ϵ {G1, G2}

~~~
d <- **dmDSdata**(counts = txiT, samples = samples)
d <- **dmFilter**(d, min_samps_feature_expr=3, min_feature_expr=5,
min_samps_gene_expr=3, min_gene_expr=20)
design_full <- model.matrix(∼Group, data=samples)
d <- **dmPrecision**(d, design=design_full)
d <- **dmFit**(d, design=design_full)
d <- **dmTest**(d, coef=“GroupG2”)
~~~

